# Differential DNA methylation in Pacific oyster reproductive tissue in response to ocean acidification

**DOI:** 10.1101/2022.03.07.483338

**Authors:** Yaamini R. Venkataraman, Samuel J. White, Steven B. Roberts

## Abstract

**Background:** There is a need to investigate mechanisms of phenotypic plasticity in marine invertebrates as negative effects of climate change, like ocean acidification, are experienced by coastal ecosystems. Environmentally-induced changes to the methylome may regulate gene expression, but methylome responses can be species- and tissue-specific. Tissue-specificity has implications for gonad tissue, as gonad-specific methylation patterns may be inherited by offspring. We used the Pacific oyster (*Crassostrea gigas)* — a model for understanding pH impacts on bivalve molecular physiology due to its genomic resources and importance in global aquaculture— to assess how low pH could impact the gonad methylome. Oysters were exposed to either low pH (7.31 ± 0.02) or ambient pH (7.82 ± 0.02) conditions for seven weeks. Whole genome bisulfite sequencing was used to identify methylated regions in female oyster gonad samples. C->T single nucleotide polymorphisms were identified and removed to ensure accurate methylation characterization.

**Results:** Analysis of gonad methylomes revealed a total of 1,284 differentially methylated loci (DML) found primarily in genes, with several genes containing multiple DML. Gene ontologies for genes containing DML were involved in development and stress response, suggesting methylation may promote gonad growth homeostasis in low pH conditions. Additionally, several of these genes were associated with cytoskeletal structure regulation, metabolism, and protein ubiquitination — commonly-observed responses to ocean acidification. Comparison of these DML with other *Crassostrea* spp. exposed to ocean acidification demonstrates that similar pathways, but not identical genes, are impacted by methylation.

**Conclusions:** Our work suggests DNA methylation may have a regulatory role in gonad and larval development, which would shape adult and offspring responses to low pH stress. Combined with existing molluscan methylome research, our work further supports the need for tissue- and species-specific studies to understand the potential regulatory role of DNA methylation.

## Background

There is great interest in elucidating how changes in the environment can impact marine invertebrate stress responses by examining DNA methylation [1, 2]. Environmental variation can alter the positions of methyl groups on CpG dinucleotides, potentially regulating gene expression by fine-tuning expression of housekeeping genes or providing additional transcriptional opportunities for environmental response genes [3]. Methylation regulation of gene expression could then influence plasticity in response to environmental stressors [1]. This framework can be applied towards understanding rapid acclimatization of invasive species [4, 5], identification of target epialleles to aid assisted evolution [6], and rearing practices for production of environmentally-resilient aquaculture species [7]. Ocean acidification [8–13], heat stress [14], upwelling conditions [15], disease [16], salinity (Johnson et al. 2021), pesticide exposure [17], and temperature and salinity [13, 18] have all elicited changes in marine invertebrate methylomes, demonstrating a potential for methylation to regulate organismal responses to a variety of environmental conditions.

Given the impact of ocean acidification on calcifying species like oysters [19–21], several environmental epigenetic studies have examined the influence of ocean acidification on molluscan methylomes (Table 1). For example, eastern oyster (*Crassostrea virginica*) mantle [10], *C. virginica* reproductive tissue [12], Hong kong oyster (*Crassostrea hongkongensis*) mantle [8], and *C. hongkongensis* larval methylomes [11] all display changes in DNA methylation after experimental ocean acidification exposure. Between all four studies, only Venkataraman et al. (2020) reports the percent of CpGs methylated: the stated 22% is consistent with the baseline 15% methylation observed in initial Pacific oyster (*Crassostrea gigas*) methylation studies [22], as well as examination of methylome responses to heat stress [14] and diuron exposure [17]. In *C. virginica*, the mantle methylome was hypomethylated after ocean acidification exposure [10], while there were no differences in global gonad methylation patterns [12]. Experimental ocean acidification exposure yielded differential methylation predominantly in gene bodies, but concentration in exons [11, 12] versus introns [8] varies. Hyper- and hypomethylation occurs with equal frequency in response to ocean acidification. This is consistent with diuron exposure [17], but not with heat stress [14]. Gene functions enriched in differential methylation datasets were inconsistent between species and tissue types (Table 1). While aminotransferase complex and biosynthetic processes were found to be enriched in hypomethylated DML in *C. virginica* mantle tissue [10], there were no enriched GOterms associated with genic DML in *C. virginica* gonad samples. In contrast, several processes were enriched in methylation datasets for *C. hongkongensis*, including acetoacetylco-A reductase activity, dehydrogenase activity, cellular response to pH, protein xylosyltransferase activity, translation factor activity, RNA binding, and diacylglycerol kinase activity in the mantle [8] and cytoskeletal and signal transduction, oxidative stress, metabolic processes, and larval metamorphosis in larvae [11]. While considerable effort has been made to understand molluscan methylation responses to ocean acidification, it is clear that there are likely species- and tissue-specific responses.

In addition to being an important global aquaculture species, *C. gigas* has been used to further our understanding of methylation in marine invertebrates [3, 7, 22–29]. Examination of diuron exposure [17] and heat stress on different *C. gigas* phenotypes [14] reveal that environmental conditions do influence the Pacific oyster methylome, and other oyster methylomes are responsive to ocean acidification (Table 1). However, studies examining the influence of ocean acidification on the *C. gigas* methylome are currently absent. Additionally, a recent study found that female *C. gigas* exposure to low pH conditions, followed by four months in ambient pH conditions prior to spawning, can still negatively impact larval survival 18 hours post-fertilization [30]. Since maturation stage was not different between female *C. gigas* after low pH exposure, this finding suggests environmental “memory” may be mediating the carryover effect [30].

Negative carryover effects were also found after parental exposure to low pH during reproductive conditioning in the hard clam *Mercenaria mercenaria* and bay scallop *Argopectan irradians* [31]. Although methylation differences associated with carryover effects have not been reported in bivalves following low pH treatment, adult *C. gigas* exposure to the pesticide diuron altered spat methylomes, demonstrating the potential for DNA methylation to influence carryover effects in bivalves [17].

We sought to understand impacts of ocean acidification on the Pacific oyster gonad methylome in order to not only understand DNA methylation as a potential mechanism for the observed negative maternal carryover effect previously reported in Venkataraman et al. [30], but to also examine the functional role of methylation in regulating biological responses to ocean acidification. Female oyster gonad methylome sequencing revealed distinct methylation changes associated with low pH exposure. These results provide the foundation for examining phenotypic plasticity within a generation, as well as larval methylomes and potential intergenerational effects.

**Table 1**. Studies exploring methylation responses to environmental stress in *Crassostrea* spp.

## Results

### Sequence Alignment

Whole genome bisulfite sequencing produced 1.38 × 10^9^ total 150 bp paired-end reads for eight total libraries. After quality trimming, 1.35 × 10^9^ total paired reads remained (<u>Supplementary Table 1</u>). Of the trimmed paired-end reads, 8.31 × 10^8^ (61.7%) were uniquely aligned to the bisulfite-converted *C. gigas* genome with appended mitochondrial sequence (<u>Supplementary Table 1</u>).

### SNP Identification

C->T SNP variants were identified from WGBS data to remove loci that might have been incorrectly characterized across individuals. A total of 13,234,183 unique SNPs were found in individual or merged BAM files, including 300,278 C->T SNPs. These C->T SNPs were used to exclude loci from downstream analyses (see DML Characterization and Enrichment Analysis).

### Global Methylation Patterns

Combined data with 5x coverage threshold provided 11,238,223 CpG loci (84.8% of 13,246,547 total CpGs in *C. gigas* genome). After C->T SNP removal, 10,939,918 CpG loci (82.6% of total *C. gigas* CpGs) were characterized. The majority of CpGs, 9,018,512 loci (82.4%), were lowly methylated, followed by 966,655 highly methylated (8.8%) and 954,751 moderately methylated (8.7%) CpGs. Highly methylated CpGs were found primarily in genic regions (903,933 CpGs, 93.5%), with 53,951 (6.0%) in exon UTR, 329,117 (36.4%) in CDS, and 523,386 (7.9%) in introns (**Figure 1**). The distribution of highly methylated CpGs was significantly different than all 5x CpGs detected by WGBS (<u>Supplementary Table 2</u>). A Principal Component Analysis (PCA) did not demonstrate any clear separation of samples by treatment (<u>Supplementary Figure 1</u>), and consistently high Pairwise Pearson’s correlation coefficients support the lack of global percent methylation differences between pH treatments (<u>Supplementary Figure 2</u>).

**Figure 1.**
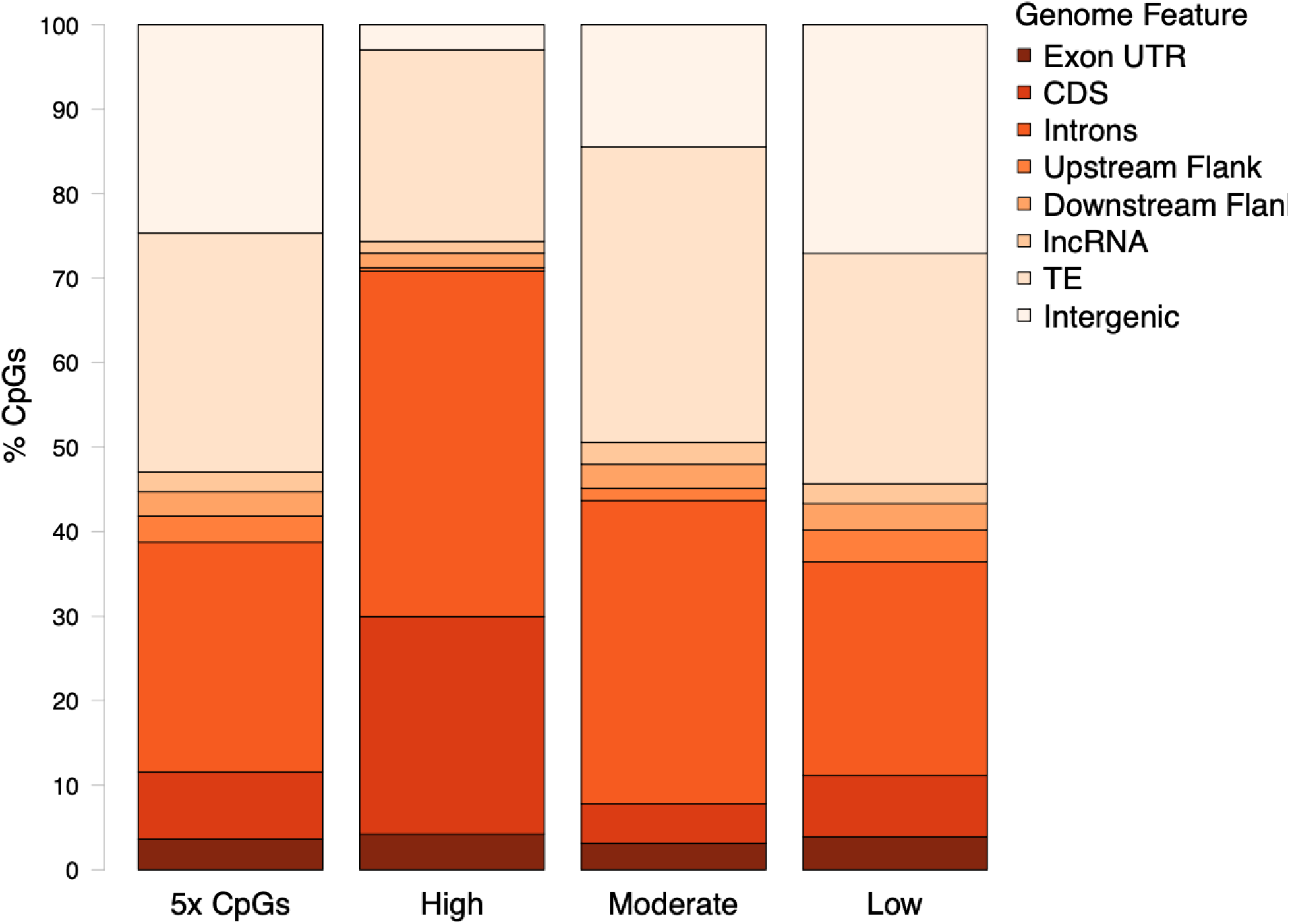
Location of all CpGs with 5x coverage. Percentage of highly methylated (≥ 50%), moderately methylated (10-50%), and lowly methylated (≤ 10%) CpGs in various genome features.

### DML Characterization

Initially, 1,599 CpG loci were identified as potential DML using a 50% methylation difference threshold. Of these CpG loci, 315 (19.7%) overlapped with C->T SNPs, and thus were removed from downstream analysis. A total of 1,284 DML were used in all remaining analyses, which consisted of 654 (50.9%) hypomethylated and 630 (49.1%) hypermethylated DML (**Figure 2**). The DML were distributed amongst the ten main chromosomes, as well as scaffolds not placed in any chromosome (**Figure 3**, <u>Supplementary Table 3</u>).

**Figure 2.**
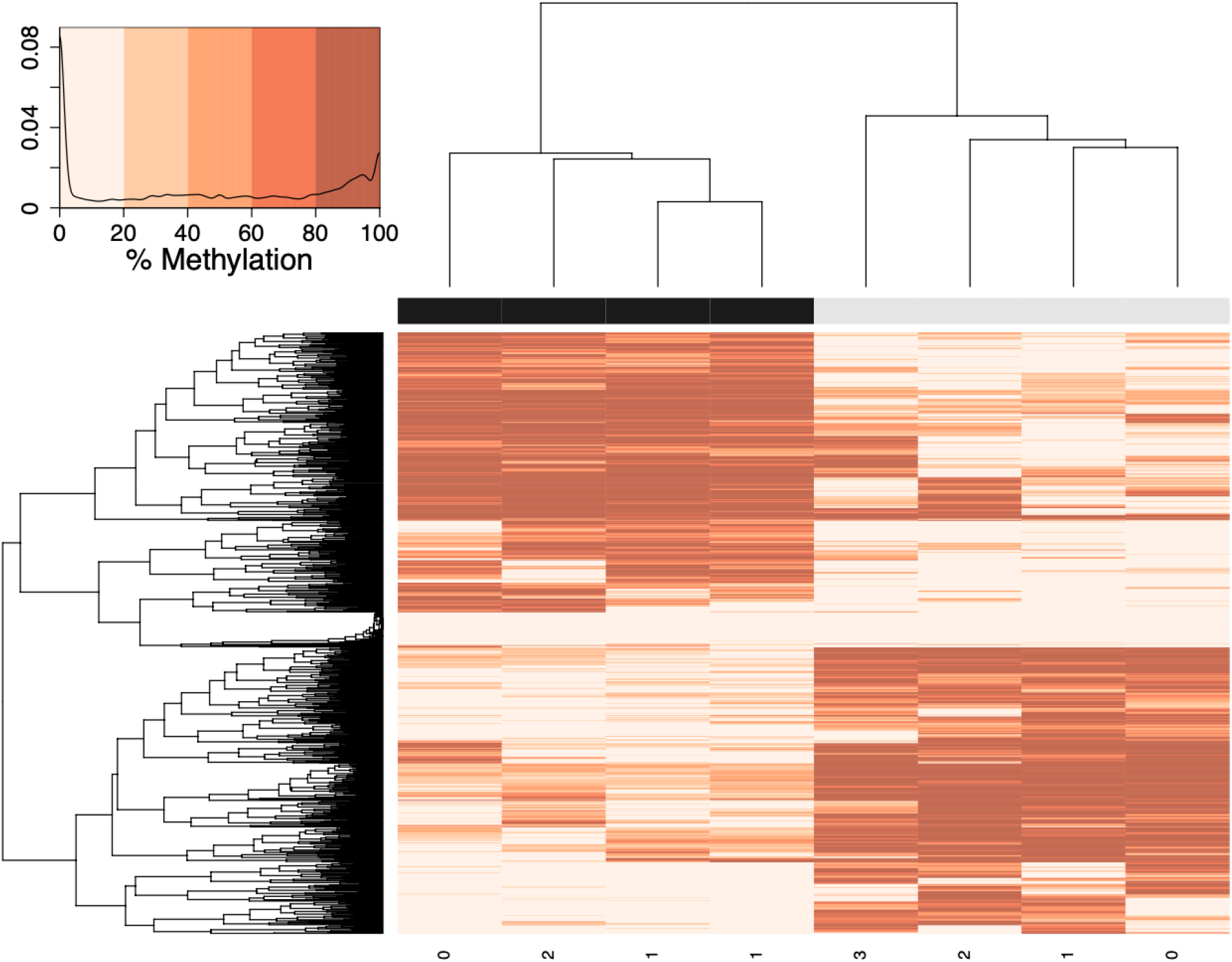
Percent methylation values for DML created using an euclidean distance matrix. Samples in low pH conditions are represented by black, and samples in ambient pH conditions are represented by gray, with maturation stage along the bottom (0 = indeterminate, 3 = spawn-ready mature female). Darker colors indicate higher percent methylation, and a density plot depicts the distribution of percent methylation values for a panel. After excluding C->T SNPs, 1,284 DML were identified using a logistic regression, using a chi-squared test and 50% methylation difference cut-off.

**Figure 3.**
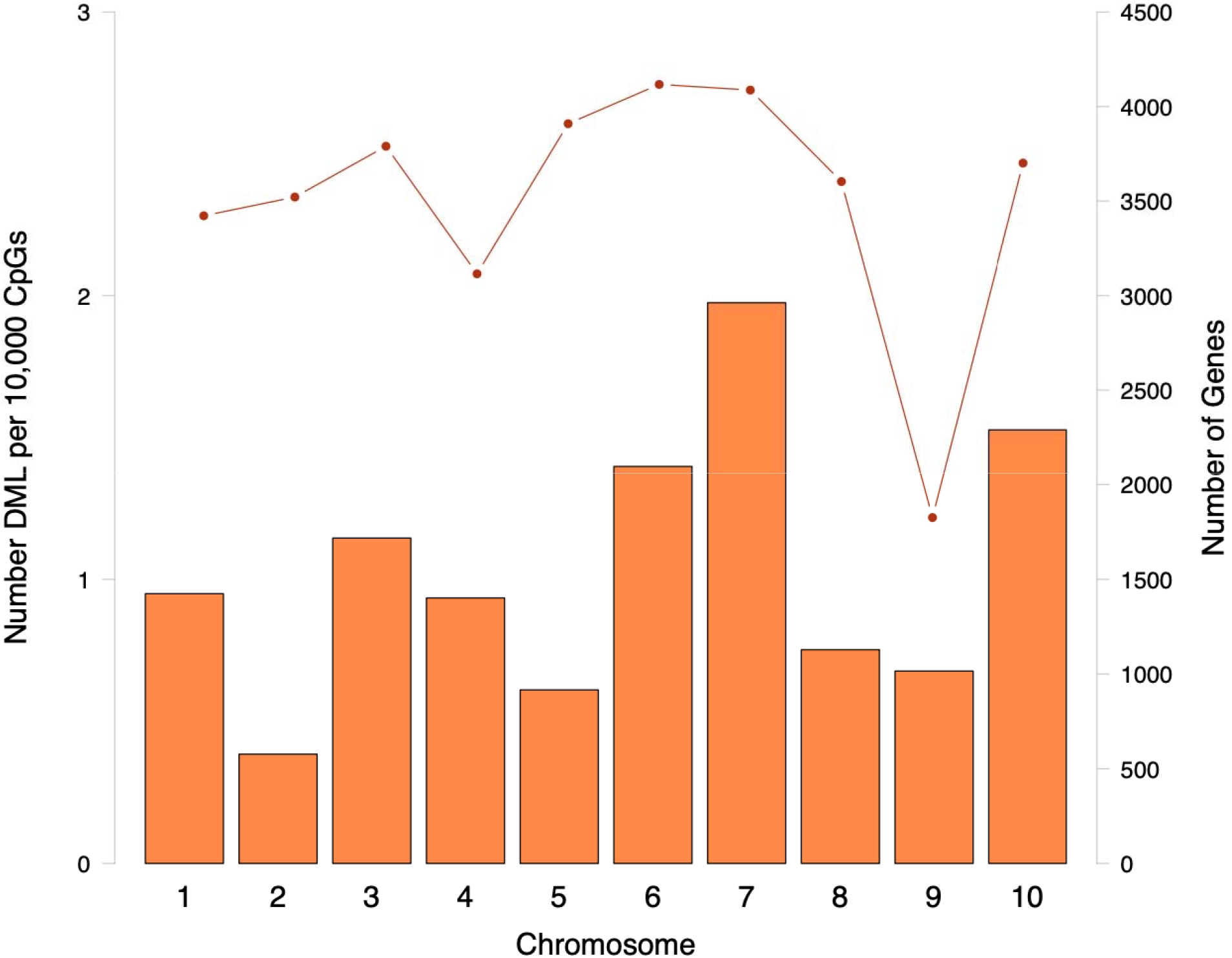
Distribution of DML in main chromosomes. Number of DML normalized by number of CpG in each chromosome (bars) and number of genes (line) in each chromosome. Additional DML were identified in scaffolds that were not mapped to any of the ten main linkage groups (<u>Supplementary Table 3</u>).

The majority of DML (1,181; 92.0%) were located in genic regions (**Figure 4**), consisting of 859 unique genes. Most genes only contained one DML; however, several genes contained multiple DML, ranging from 2-103 DML in one gene (<u>Supplementary Table 4</u>). The genic DML were primarily located in introns (783; 61.0%), followed by CDS (442; 34.4%) and exon UTR (77; 6.0%). Putative promoter regions upstream of transcription start sites contained DML (10; 0.8%), but more were located in downstream flanking regions (61; 4.8%). The DML were also found in transposable elements (434; 33.8%), lncRNA (9; 0.7%), and intergenic regions (38; 3.0%). The number of DML in all genome features differed significantly from all 5x CpGs (<u>Supplementary Table 5</u>).

**Figure 4.**
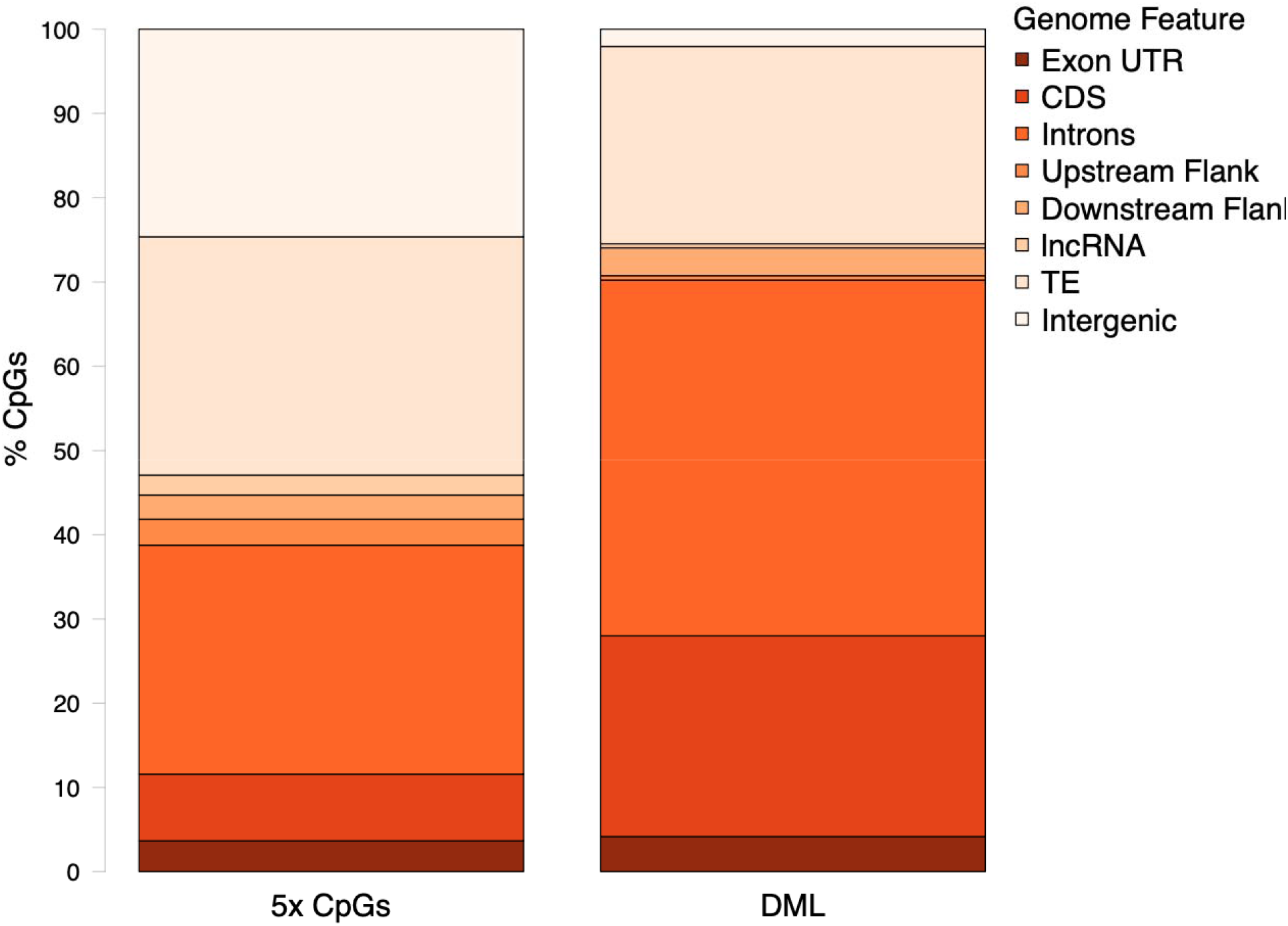
Location of CpGs with 5x coverage and DML. Percentage of 5x CpGs and DML found in various genome features.

### Enrichment Analysis

Ten biological processes and 4 cellular component GOterms were significantly enriched in genes containing DML. A total of 339 genes with GOterm annotations containing 437 DML were used for this analysis. The enriched biological process GOterms were involved in motility (ex. cilium-dependent cell motility, cilium movement) and development (ex. insta larval or pupal development, post-embryonic development) (<u>Supplementary Table 6</u>). Nine unique genes (37 associated transcripts) contained enriched biological process GOterms: *dynein heavy chain 5, axonemal, unconventional mysoin-VI, serine/threonine-protein kinase 36, helicase domino, protein neuralized, cytoplasmic aconitate hydratase, serine-protein kinase ATM, cation-independent mannose-6-phosphase receptor, uncharacterized LOC105324167*. These genes contained 16 DML, 13 of which were hypermethylated (<u>Supplementary Table 6</u>). Enriched cellular component GOterms were involved in microtubule-associated complexes and various acetyltransferase complexes (<u>Supplementary Table 7</u>). The six unique genes (25 associated transcripts) that contained enriched cellular component GOterms were *dynein heavy chain 5, axonemal, cytoplasmic dynein 2 light intermediate chain-1-like, unconventional myosin-VI, helicase domino, and kinesin-like protein KIF23* (<u>Supplementary Table 7</u>). Six of the seven DML contained in these genes were hypermethylated (<u>Supplementary Table 7</u>).

## Discussion

Our work examined environmentally-induced changes to the *C. gigas* reproductive tissue methylome. We identified a total of 1,284 differentially methylated loci, showing that the methylome is responsive to ocean acidification. We also found enriched biological processes associated with genic DML, implying that low pH may impact regulation of distinct processes, such as gonad development. Taken together, our findings suggest that environmentally-responsive methylation may maintain gonad development in stressful conditions. Our findings contribute to the growing body of work examining epigenetic responses in molluscan species, and provide the foundation for future research examining intergenerational epigenetic inheritance and its potential to mediate plastic responses to environmental stressors.

### Functions of genes containing DML with enriched GOterms suggest role for methylation regulation of gonad development

Enrichment of microtubule associated complex and cilium or flagellar motility GOterms imply methylation may control microtubule movement in response to low pH. Motor proteins like *dynein heavy chain 5, axonemal*, and *cytoplasmic dynein 2 light intermediate chain 1-like* form complexes to move flagella or cilia. Dyneins exhibit higher expression over the course of *C. gigas* spermatogenesis [32]. In female gonad tissue, hypermethylation may suppress expression of dyneins to promote oogenesis. Indeed, lack of significant sex ratio and maturation stage differences between low and ambient pH treatments suggests that oogenesis was maintained in low pH conditions [30]. Studies examining molluscan somatic tissues have found low pH impacts on dynein expression [33–35], with significantly lower dynein protein abundance [33] or gene expression [34] in oysters exposed to ocean acidification. Similar to cytoplasmic dyneins, *unconventional myosin-VI* plays a role in organelle movement. Myosins generally exhibit high expression in immature gonads, which reduces as the gonad matures [32, 36]. The reduction in myosin expression may signify reorganization of female reproductive tissue to accommodate growing oocytes. Hyper- and hypomethylation of unconventional myosin may regulate cytoskeletal structure of cells to promote growth of reproductive over somatic tissue. Dynein and myosin genes exhibit differential methylation in response to low pH in *C. virginica* gonad as well [12]. Regulation of motility genes may be a conserved response to ocean acidification, and methylation of these genes may promote homeostasis during gametogenesis.

Several genes with DML and enriched GOterms have links to stress response and gonad development. In *C. gigas*, expression of cell cycle regulators *kinesin-like protein KIF23* and *serine-protein kinase ATM* changes in response to immune challenge or low pH and arsenic exposure, respectively [37, 38]. Kinesin-like proteins have higher expression in mature *C. gigas* gonads [32], and serine-protein kinase ATM is involved in razor clam *Sinonovacula constricta* oogenesis [39]. Another kinase, *serine/threonine-protein kinase 36*, is involved in signaling pathways. Several serine/threonine-protein kinases are highly expressed in mature *C. gigas* female gonad, with certain homologues more expressed in females with mature gonad and during sex determination and differentiation events [32, 36]. Serine/threonine-protein kinases are also related to oocyte maturation in the king scallop *Pecten maximus* [40]. An investigation of the *C. gigas* kinome found various serine/threonine-protein kinases present in eggs, including *serine/threonine-protein kinase 36*, that were responsive to abiotic stressors [41].

Serine/threonine-protein kinase genes were also differentially methylated in *C. virginica* gonad [12]. As a catalyst of the TCA cycle, *cytoplasmic aconitate hydratase* is upregulated during spermatogenesis in *C. gigas* and downregulated during spermatogenesis in *P. maximus*, with differing consequences for energy metabolism [42, 43]. Early examination of *C. virginia* found *cytoplasmic aconitate hydratase* localized to eggs, with enzyme activity increasing during embryogenesis [44]. Therefore, it’s possible *cytoplasmic aconitate hydratase* may regulate energy metabolism during oogenesis as well. Evidence of *cytoplasmic aconitate hydratase* abundance changing in response to low pH has been documented in *C. gigas*, which implies that low pH alters energy metabolism [38]. Energy metabolism would be important to regulate in developing oocytes. *Helicase domino* regulates transcription through histone phosphorylation, acetylation, and chromatin remodeling. Populations of the Olympia oyster (*Ostrea lurida*) contain an outlier SNP loci in the *helicase domino* gene, which is implicated in immune or stress response [45]. Upregulation of *helicase domino* was associated with oogenesis maintenance in *Acartia tonsa* shrimp [46]. Taken together, hypermethylation of *kinesin-like protein 23, serine-protein kinase ATM, cytoplasmic aconite hydratase*, and *helicase domino*, and hypomethylation of *serine/threonine-protein kinase 36*, suggests that methylation may regulate gonad development in stressful conditions, allowing the oyster to maintain homeostasis.

### Similarities between differential methylation in gonads and larvae

Investigation of *C. hongkongensis* larvae exposed to ocean acidification revealed enrichment of cytoskeletal and signal transduction processes in differentially methylated genes [11]. Genes with DML in *C. gigas* gonad are enriched for similar processes, and have links to embryonic development. Unconventional myosins, important cytoskeletal components, were found to be important for fertilization, blastulation, and gastrulation in *S. purpuratus* [47]. Ocean acidification drove decreased expression of another cytoskeletal component, kinesin-like protein, in *S. purpuratus* larvae, which was related to measured reductions in calcification [48]. Expression of signal transduction kinases found in embryos changed in response to various abiotic stressors [41]. *Serine/threonine protein-kinase pim-3* gene was differentially methylated in *C. hongkongensis* larvae [11], and this study found a DML in *serine/threonine-protein kinase 36* in *C. gigas* gonad. Differential methylation in cytoskeleton and signal transduction genes in female gonad may influence larval responses to low pH.

Enrichment of metabolic processes in *C. gigas* gonad and *C. hongkongensis* larval methylation datasets [11] signify that methylation regulation of metabolism is important across life stages. In *S. purpuratus* with a history of low pH exposure, day-old embryos exhibited higher expression of *cytoplasmic aconitate hydratase* than embryos from populations without exposure history [49]. However, larvae from populations without exposure history had higher expression of *cytoplasmic aconitate hydratase* after seven days [49]. Modulation of the TCA cycle through *cytoplasmic aconitate hydratase* may be important for metabolic regulation in developing larvae, and inheritance of hypermethylation within this gene could prime larval metabolism for low pH exposure. *Cation-independent mannose-6-phosphate receptor* is a lysosomal enzyme associated with post-embryonic development (GO:0009791). Expression of this gene was upregulated during zebra mussel *Dreissena polymorpha* and shrimp *Marsupenaus japonicus* immune challenges [50, 51]. Although no current research has explored this gene’s role in larval development or abiotic stress response, hypomethylation of *cation-independent mannose-6-phosphate receptor* may impact lysosome metabolic activity and immune function during a critical growth period. *Protein neuralized* is part of a group of ubiquitin ligases. Regulation of ubiquitination is likely a conserved response to ocean acidification in oysters: elevated pCO_2_ increased abundance of proteins involved in ubiquitination and decreased protein degradation in *C. gigas* posterior gill lamellae [33], and upregulated ubiquitination gene expression in adult *C. virginica* mantle [52]. Differential methylation of protein ubiquitination genes was also found in *C. virginica* gonad [12] and *C. hongkongensis* larvae [11]. Hypermethylated DML within the *protein neuralized* gene could regulate ubiquitination. Differential methylation in this gene may also regulate cell division and larval growth, as upregulation of protein neuralized has been found to suppress cell division and macromolecule synthesis in *Artemia sinica* during diapause [53].

Taken together, methylome similarities between *C. gigas* female gonad and *C. hongkongensis* larvae suggest that differential methylation in these genes could also impact larval calcification, cellular structure, and metabolism if patterns are inherited. Intergenerational methylation inheritance has been demonstrated previously in *C. gigas*: adult exposure to the pesticide diuron altered spat methylomes [17]. While we did not investigate *C. gigas* larval methylomes, we did find that prior exposure of female oysters to low pH conditions negatively impacted larval survival in ambient pH conditions 18 hours after fertilization [30]. It is possible that the methylation patterns responsible for maintaining homeostasis in female gonad tissue were maladaptive in larvae, especially when parent and larval environments were mismatched.

### Considerations for marine invertebrate methylation research

The concentration of DML in genes is consistent with the literature examining molluscan methylome responses to ocean acidification [8, 10–12], disease [16], pesticide exposure [17], salinity (Johnson et al. 2021), and heat stress [14], as well as environmental responses in corals [13, 18], urchins [15], and pteropods [9]. This differs from the vertebrate model of methylation, in which CpG islands are found in promoter regions upstream of transcription start sites [1]. Similar pathways exhibiting differential methylation — but not exact genes — between *C. gigas* gonad (this study), *C. virginica* gonad [12] and mantle [10], and *C. hongkongensis* mantle [8] and larvae [11] suggest tissue- and species-specific methylome responses to low pH. In *C. gigas*, high levels of gene methylation are associated with increased gene expression and low expression variability, implying that methylation leads to transcriptional stability [22, 25, 29].

Similarly, exposure to the pesticide diuron modified the *C. gigas* methylome such that DNA methylation changes correlated with RNA abundance in a small group of highly methylated genes [17], and increased methylation was also found to reduce transcriptional noise in response to ocean acidification in corals [13]. While recent *Crassostrea* spp. studies have found no connection between differentially methylated and differentially expressed genes after exposure to low pH [8, 10], individual DML may play a role in shaping gene expression.

Comparison of a single gene with differing methylation levels between *C. gigas* mantle and male gametes demonstrated that gene methylation was associated with alternative splicing and gene expression [29]. Therefore, it is critical to examine how individual DML can influence gene activity in response to environmental change.

The role of DML in shaping gene activity is likely dependent on their locations in genome features. We found 442 and 941 DML in introns and CDS, respectively, with roughly half of the genic DML being hypermethylated in treatment gonads. In *C. gigas*, exons included in the final transcript and longer exons generally have higher methylation [29]. Genes with higher methylation in exons or introns were highly expressed in *S. purpuratus*, and had a higher likelihood of alternative splicing, alternative start site, or exon skipping in response to upwelling conditions [54]. Differentially methylated positions in introns and CDS may have similar roles in *C. gigas* response to low pH. Additionally, presence of multiple DML in a gene may have conflicting effects on expression. Most genes examined in this study only contained one DML; however, we found several genes with multiple DML, ranging from 2-103 in one gene. High methylation in both exons and introns did not produce high expression levels in *S. purpuratus*, suggesting increased methylation in both of these genome features act antagonistically to impact gene expression [54]. Multiple DML in CDS and introns may influence *C. gigas* expression similarly. Future work should examine how environmentally-influenced methylation impacts gene expression by considering individual DML, not averaged methylation for a gene.

Confounding elements may have influenced gonad methylome responses to ocean acidification, and our interpretation of those responses. Maturation stage is known to produce distinct baseline methylation patterns in *C. gigas* (Zhang et al., 2018). While the overall maturation stage was used as a covariate when identifying DML, it is possible that each sample methylome contains information from cells with various maturation stages or cell types. Single-cell sequencing efforts or investigations within a single maturation stage would clarify how these sources of variation influenced our findings. We also identified several C->T SNPs that overlapped with DML even though specimens were sibling oysters with similar genomic backgrounds. Recent examination of *C. virginica* methylomes suggest methylation is largely linked to genotype, emphasizing the need to account for genotypic differences in methylation calls [55]. Our work demonstrates how SNP identification from WGBS can be used to exclude loci with incorrectly characterized methylation due to genomic variation. While the lack of annotated genes used in the enrichment analysis limits our interpretation of which processes are most impacted by pH-sensitive methylation, removing SNP-DML overlaps prior to enrichment analysis provides a more biologically accurate understanding of how methylation may impact gonad growth.

## Conclusion

Our work — which is the first to explore DNA methylation responses to ocean acidification in Pacific oysters — found that low pH treatment altered the *C. gigas* female gonad methylome after a seven-week exposure. We found that differential methylation occurs primarily in genic regions, which is consistent with other studies examining oyster methylome responses to ocean acidification. In conjunction with histology data from Venkataraman et al. [30], enriched biological processes and cellular components in genes containing DML suggest that methylation may be used to maintain gonad growth in low pH conditions. Our work expands on previous molluscan methylation research by not only demonstrating a potential role for methylation in response to ocean acidification, but also showing the species- and tissue-specific nature of these responses.

As inheritance of environmentally-induced methylation — and any resulting phenotypic plasticity — is contingent on epigenetic marks being present in the germline, withstanding chromatin reorganization, and being packaged into gametes [1], the next step in this work is correlating germline methylation with offspring methylation, and examining gene expression, alternative splicing, chromatin regulation, and phenotype. Comparing parental and offspring phenotype will allow researchers to assess whether parental experience and resulting epigenetic responses are accurate, reliable cues for adaptive plasticity in offspring.

## Methods

### DNA Extraction and Library Preparation

Adult hatchery-raised *C. gigas* (average shell length = 117.46 ± 19.16 mm) were exposed to either low pH (7.31 ± 0.02) or ambient pH (7.82 ± 0.02) conditions from February 15, 2017 to April 8, 2017 at the Kenneth K. Chew Center for Shellfish Research and Restoration at the National Oceanic and Atmospheric Administration Manchester Field Station (47º 34’ 09.1” N 122º 33’ 19.0” W, Manchester, WA). The experimental design, seawater chemistry analysis with carbon chemistry parameters, and histological analysis are described in Venkataraman et al. [30]. To determine how pH exposure altered maternal gonad methylation, four female (or presumed female) oysters sampled at the end of the seven-week pH exposure were selected for each treatment. DNA was extracted from histology blocks for these gonad tissue samples using the PAXgene Tissue DNA Kit (Qiagen) with modifications specified below. For each sample, up to 0.02 g of histology tissue embedded in paraffin was cut from the histology block and placed in a 2 mL round-bottomed tube. The tissue was then further homogenized within the tube. After adding 1 mL of xylene, each sample was vortexed for 20 seconds, incubated at room temperature for three minutes, then centrifuged at maximum speed for three minutes. To evaporate ethanol from samples after ethanol addition and vortexing steps, samples were placed on a heat block at 37ºC for 10-20 minutes.

The resuspended pellet was lysed using a TissueTearor at setting 1. Prior to lysis and between samples, the TissueTearor was run in a 10% bleach solution for 20 seconds, followed by two consecutive 20 second runs in DI water to clean the instrument. Lysates were transferred to clean, labeled 1.5 mL microcentrifuge-safelock tubes. Proteinase K (20 μL) was added to each sample, and pulse-vortexed for 15 seconds to mix. The sample-Proteinase K solution was incubated 56ºC for 60 minutes. Every ten minutes, the samples were briefly removed from the heat block to vortex at maximum speed for one minute. After 60 minutes, RNase A (4 μL, 100 mg/mL) was added to each sample. Samples were vortexed at maximum speed for 20 seconds to mix and incubated for two minutes at room temperature to obtain RNA-free genomic DNA and reduce possible interference for downstream enzyme reactions. Samples were then kept at 80ºC for 60 minutes, and vortexed at maximum speed for one minute in ten-minute intervals. After the incubation, Buffer TD2 and ethanol (200 μL) were added to each sample, vortexing thoroughly to mix.

DNA was isolated for each sample after lysis and incubation following manufacturer’s instructions. To elute DNA, samples were loaded onto the spin column with Buffer TD5 (50 μL) and incubated at room temperature for five minutes to increase yield, then centrifuged at maximum speed for one minute. A Qubit™ 3.0 and dsDNA BR Assay (Invitrogen) was used to quantify sample yield using 1 μL of DNA for each sample. For two samples, DNA was further concentrated using an ethanol precipitation.

Bisulfite conversion was performed with the Zymo-Seq WGBS Library Kit (Cat. #D5465) using 100 ng of genomic DNA following manufacturer’s instructions. Samples were spiked with a mixture of six unique double-stranded synthetic amplicons (180-200 bp) from the *Escherichia coli* K12 genome. Since each amplicon represents a different percent methylation, the spike-in was used to determine bisulfite conversion efficiency. Libraries were barcoded and pooled into a single lane to generate 150 bp paired-end reads in a single Illumina NovaSeq flowcell.

### Genome Information and Feature Tracks

Nuclear genome [56] (NCBI GenBank: GCA_902806645.1, RefSeq: GCF_902806645.1) and mitochondrial genome sequences (NCBI Reference Sequence: NC_001276.1) were used in downstream analysis. These sequences were combined for downstream alignment. The `fuzznuc` function within EMBOSS was used to identify CG motifs in the combined genomes.

The RefSeq annotation was used to obtain genome feature tracks, and create additional tracks. The annotation included a combination of Gnomon, RefSeq, cmsearch, and tRNAscan-SE predictions. Gene, coding sequence, exon, and lncRNA tracks were extracted from the RefSeq annotation. These tracks were then used to obtain additional genome feature information. To create an intron track, the complement of the exon track was generated with BEDtools `complementBed` v.2.26.0 to create a non-coding sequence track [57]. The overlap between the genes and the non-coding sequences was found with `intersectBed` and defined as introns.

Similarly, untranslated regions of exons were obtained by subtracting coding sequences from exon information with `subtractBed`. Flanking regions were defined as the 1000 bp upstream or downstream of genes. Upstream flanks were generated by adding 1000 bp upstream of genes, taking into account strand, with `flankBed`. Existing genes were removed from flanking sequences using `subtractBed`. A similar process was performed to generate downstream flanking regions. Intergenic regions were isolated by finding the complement of genes with `complementBed`, then using `subtractBed` to remove any flanks. Transposable element locations were obtained from RepeatMasker using the NCBI RefSeq annotation (RefSeq: GCF_902806645.1) [58, 59]. The number of CG motifs in a given feature track was obtained using `intersectBed`. All genome feature tracks are available in the associated Open Science Framework repository (see *Availability of data and materials*).

### Sequence Alignment

Reads were trimmed three times prior to alignment using TrimGalore! v.0.6.6 [60]. Known Illumina adapter sequences were removed (`--illumina`), along with 10 bp from both the 5’ (`--clip_R1 10` and `--clip_R2 10`) and 3’ ends (`--three_prime_clip_R1 10` and `--three_prime_clip_R2 10`) of 150 bp paired-end reads. Any remaining adapters were removed in a second round of trimming. Finally, poly-G tails were trimmed out of samples by manually specifying adapter sequences (`--adapter GGGGGGGGGGGGGGGGGGGGGGGGGGGGGGGGGGGGGGGGGGGGGGGGGG` and `--adapter2 GGGGGGGGGGGGGGGGGGGGGGGGGGGGGGGGGGGGGGGGGGGGGGGGGG`). Sequence quality was assessed with FastQC v.0.11.9 [61] and MultiQC [62] after each round of trimming.

Trimmed samples were then aligned to the combined nuclear and mitochondrial genomes for *C. gigas* [56]. The genome was bisulfite converted with Bowtie 2-2.3.4 (Linux x84_64 version [63]) using the `bismark_genome_preparation` function within Bismark v.0.22.3 [64]. Reads were aligned to the bisulfite-converted genome. Non-directional input was specified (`--non_directional`), and alignment scores could be no lower than -90 (`-score_min L,0,-0.6`). Resultant BAM files were deduplicated (`--deduplicate_bismark`), and methylation calls were extracted from deduplicated BAM files (`bismark_methylation_extractor`).

Deduplicated BAM files were also sorted and indexed for downstream applications using SAMtools v.1.10 [65]. Using coverage files generated from methylation extraction, CpG information was consolidated between strands to report 1-based chromosomal coordinates (`coverage2cystosine`). Using `coverage2cytosine` coverage file output, genomic coordinates for CpGs with at least 5x coverage across all samples were written to BEDgraphs. Summary reports with alignment information for each sample were generated (`bismark2report` and `bismark2summary`) and concatenated with MultiQC. Code for all sequence alignment steps can be found in the associated Open Science Framework repository (see *Availability of data and materials*).

### SNP Identification

During bisulfite conversion, cytosines methylated in a CpG context are retained as cytosines, while unmethylated cytosines are converted to uracils. These uracils are then converted to thymines during PCR amplification. As C->T single nucleotide polymorphisms (SNPs) would be interpreted by Bismark as an unmethylated cytosine, removing these loci is an important step to ensure accurate methylation characterization. SNP variants across all samples and in individuals were identified with `BS-Snper` [66]. This program takes into consideration nucleotide identity on the opposite strand: a thymine paired with an adenine indicates a C->T SNP, while a thymine paired with a guanine is a bisulfite-converted unmethylated cytosine. Sorted deduplicated BAM files generated with Bismark v.0.22.3 and SAMtools v.1.10 [64, 65] were merged (`samtools merge`) to identify SNPs. Variants were also identified in individual sorted deduplicated BAM files. Default `BS-Snper` settings were used, except for a minimum coverage of 5x to be consistent with methylation analyses. The C->T SNPs were used to exclude loci of interest.

### Global Methylation Characterization

Global methylation patterns were characterized by averaging DNA methylation across all samples using loci with at least 5x coverage that did not overlap with C->T SNPs. A CpG locus was considered highly methylated if average percent methylation was greater than or equal to 50%, moderately methylated if percent methylation was between 10-50%, and lowly methylated if percent methylation was less than or equal to 10%. The genomic location of highly, moderately, and lowly methylated CpGs in relation to UTR, CDS, introns, up- and downstream flanking regions, transposable elements, and intergenic regions were determined by using `intersectBed`. We tested the null hypothesis that there was no association between the genomic location of CpG loci and methylation status (all CpGs versus highly methylated CpGs) with a chi-squared contingency test (`chisq.test` in R Version 3.5.3 [67]).

### Identification of Differentially Methylated Loci

Identification of differentially methylated loci, or DML, was conducted with `methylKit v.1.17.4` [68] in R Version 3.5.3 [67]. Prior to DML identification, data were imported and processed. Data with at least 2x coverage were imported using `methRead` from merged CpG coverage files produced by `coverage2cytosine`. Imported data were filtered again (`filterByCoverage`) to require 5x coverage for each CpG locus (`lo.count = 5`) in each sample. Potential PCR duplicates in each sample were also removed by excluding CpG loci in the 99.9th percentile of coverage (`high.perc = 99.9`). Once filtered, data were normalized across samples (`normalizeCoverage`) to avoid over-sampling reads from one sample during downstream statistical analysis. Histograms of percent methylation (`getMethylationStats`) and CpG coverage (`getCoverageStats`) were used to confirm normalization. After filtering and normalization, data at each CpG locus with 5x were combined (`unite`) to create a CpG background, ensuring that each locus had at least 5x coverage in each sample.

Methylation differences were then identified for *C. gigas* gonad samples. A logistic regression (`calculateDiffMeth`) modeled the log odds ratio based on the proportion of methylation at each CpG locus. A covariate matrix with gonad stage information (`covariates`) was applied to the model for the *C. gigas* data to account for varying maturation stages in samples. Additionally, an overdispersion correction with a chi-squared test (`overdispersion = “MN”, test = “Chisq”`) was applied.

Differentially methylated loci, or DML, were defined as CpG dinucleotides that did not overlap with C->T SNPs with at least a 50% methylation difference between pH treatments, and a q-value < 0.01 based on correction for false discovery rate with the SLIM method [69]. Hypermethylated DML were defined as those with significantly higher percent methylation in low pH samples, and hypomethylated DML with significantly lower percent methylation in low pH samples. BEDfiles with DML locations were created and viewed with the Integrative Genomics Viewer (IGV) version 2.9.2 [70].

### DML Characterization

The location of DML was characterized in relation to various genome feature tracks. Presence of DML in UTR, CDS, introns, up- and downstream flanking regions, transposable elements, and intergenic regions was discerned using `intersectBed`. A chi-squared contingency test was used to test the null hypothesis of no association between genomic location and differential methylation by comparing the genomic location of all CpGs with 5x data and DML. Additionally, the number of DML present in each chromosome was quantified to see if DML were distributed uniformly across the genome. Overlaps between DML and SNPs were identified to determine if DML were solely attributed to treatment differences, or if genetic differences contributed to differential methylation.

### Enrichment Analysis

To determine if genes containing DML were associated with overrepresented biological processes, cellular component, or molecular function gene ontologies (GO), functional enrichment analysis was performed. Prior to this analysis, the *C. gigas* genome was annotated. A `blastx` alignment was performed against the Uniprot-SwissProt database (accessed June 01, 2021) to get Uniprot Accession information for each transcript [71]. The transcript nucleotide sequences were extracted from the genome (GCF_902806645.1_cgigas_uk_roslin_v1_rna_from_genomic.fna available in the <u>NCBI Annotation Release 102</u>). Transcript IDs from the `blastx` output were matched with GO terms from the Uniprot-Swissprot database using Uniprot Accession codes. These transcript IDs and corresponding GO terms were then used to create a gene ID-to-GO term database (`geneID2GO`) for manual GO term annotation. The transcript IDs were filtered for genes that contained CpGs with at least 5x coverage in all samples, as the same parameters were used to generate the CpG background for differential methylation analysis. Each line of the database contained transcript ID in one column, and all corresponding GO terms in another column. Gene enrichment analysis was conducted with `topGO` v.2.34.0 in R [72]. The `geneID2GO` database was used as the gene universe for enrichment, while transcript IDs for genes with DML were used as the genes of interest. These transcript IDs were filtered such that they did not include any transcripts associated with DML-SNP overlaps. A `topGO` object was generated for each DML list and GO category (biological process, cellular component, or molecular function), with GO term annotation performed using the `geneID2GO` database. A Fisher’s exact test was used to identify GO terms in each DML list significantly enriched with respect to the gene background (*P*-value < 0.01). Keeping with `topGO` default settings, we did not correct for multiple comparisons [72]. Enriched GOterms were clustered by semantic similarity using `simplifyEnrichment` v.1.2.0 default settings [73]. A semantic similarity matrix was created from enriched GO IDs using the “Relevance” similarity measure and GO tree topology (`GO_similarity`). The semantic similarity matrix was clustered using the binary cut method, and visualized as a word cloud alongside a heatmap of semantic similarity (`simplifyGO`). Finally, the enriched GOterms and cluster information were appended to a list of genic DML and annotations to understand which genes with DML also had enriched GOterms.

## List of Abbreviations

WGBS: whole genome bisulfite sequencing
DML: differentially methylated locus/loci
SNP: single nucleotide polymorphism

## Declarations

### Ethics approval and consent to participate

Not applicable.

### Consent for publication

Not applicable.

### Availability of data and materials

All data, genome feature tracks, scripts, and a supplementary materials list are available in the Oyster Gonad Methylation repository, doi.org/10.17605/OSF.IO/YGCTB. All raw data can be accessed at the NCBI Sequence Read Archive under BioProject accession number PRJNA806944 (https://www.ncbi.nlm.nih.gov/bioproject/806944).

### Competing interests

The authors declare that they have no competing interests.

### Funding

This work was funded by National Science Foundation award 1634167 to SBR. The Hall Conservation Genetics Research Fund (YRV) supported sequencing for this project.

### Authors’ contributions

YRV conceived the experimental design and conducted the pH exposure experiment with assistance from SBR. YRV extracted DNA for sequencing and analyzed sequence data with SJW and SBR. YRV wrote the initial manuscript with input from SJW and SBR. All authors reviewed and approved the final manuscript.

## Acknowledgements

Grace Crandall performed histological analysis and Kaitlyn Mitchell assisted with DNA extractions. This work was facilitated through the use of advanced computational, storage, and networking infrastructure provided by the Hyak supercomputer system at the University of Washington.

## Notes

### Competing Interest Statement

The authors have declared no competing interest.

